# Proteomics reveals extensive phosphoregulation of outer kinetochore protein KNL1

**DOI:** 10.64898/2026.03.13.711714

**Authors:** Abby C Jurasin, Anderson R Frank, Sue Biggins

## Abstract

Microtubules attach to kinetochores to facilitate chromosome movement to opposite spindle poles. Defective kinetochore-microtubule attachments lead to phosphorylation of the outer kinetochore protein KNL1 at conserved MELT motifs, which triggers spindle assembly checkpoint activation and recruitment of the fibrous corona. To identify additional phosphorylation sites that regulate kinetochores, we treated HEK 293T/17 cells with nocodazole, paclitaxel, or STLC to create defective kinetochore-microtubule attachment states. We then purified KNL1 and performed proteomics and identified 111 phosphorylation sites on KNL1, including several that may be attachment-state specific. These data demonstrate that KNL1 is extensively phosphoregulated in response to treatment with microtubule-disrupting compounds.

## Description

Chromosome segregation relies on attachments between microtubules and kinetochores, multi-megadalton protein complexes that assemble on centromeric DNA during cell division. Proper attachments generate tension, which stabilizes the kinetochore-microtubule bond (Nicklas and Koch 1969; Nicklas and Ward 1994; King and Nicklas 2000; Akiyoshi et al., 2010). Erroneous attachments, if left uncorrected, result in chromosome mis-segregation and aneuploidy, which have been implicated in cancer development (McAinsh and Kops 2023). Cells utilize several mechanisms to combat these risks, including activation of the spindle assembly checkpoint and recruitment of an expansive protein network referred to as the fibrous corona. The spindle assembly checkpoint surveils for defective kinetochore-microtubule attachments and prevents progression into anaphase until proper attachments are established (McAinsh and Kops 2023). The fibrous corona forms a crescent-like structure at the outer kinetochore to facilitate microtubule recapture (Wynne and Funabiki 2015; Kops and Gassmann 2020).

Spindle assembly checkpoint activation and corona recruitment are both dependent on KNL1 phosphorylation. Specifically, phosphorylation of MELT and SHT motifs by the dual specificity threonine tyrosine kinase (TTK, also known as MPS1) and polo-like kinase 1 (PLK1), initiates a signaling cascade that results in production of the soluble mitotic checkpoint complex (MCC) (von Schubert et al., 2015; McAinsh and Kops 2023). The MCC then inhibits the anaphase promoting complex/cyclosome (APC/C), thus halting mitosis before anaphase onset (von Schubert et al., 2015; McAinsh and Kops 2023). In addition to its role in spindle assembly checkpoint activation, KNL1 phosphorylation also promotes oligomerization and lateral expansion of several spindle assembly checkpoint associated proteins to form the fibrous corona (Wynne and Funabiki 2015; Kops and Gassmann 2020). There are several kinetochore-microtubule attachment states that induce spindle assembly checkpoint activation, but prior studies have largely focused on unattached kinetochores. It remains unclear whether KNL1 phosphorylation, recruitment of spindle assembly checkpoint proteins, and corona expansion may differ in other erroneous kinetochore-microtubule attachment states.

To evaluate changes in KNL1 interaction partners and phosphorylation across different kinetochore-microtubule attachment states, we treated HEK 293T/17 cells with one of 3 microtubule-disrupting compounds: nocodazole, paclitaxel, or *S*-trityl-L-cysteine (STLC). Nocodazole inhibits microtubule polymerization, resulting in a complete loss of kinetochore-microtubule attachments (Fig. 1A) (Zieve et al., 1980). Paclitaxel stabilizes microtubules, restricting the dynamicity that is critical for the establishment of end-on attachments (Fig. 1A) (Schiff et al., 1979). STLC inhibits the kinesin Eg5, which is required for the separation of centrosomes and establishment of a bipolar mitotic spindle (Fig. 1A) (Skoufias et al., 2006). Following STLC treatment, sister kinetochores form syntelic attachments to microtubules emanating from this monopolar spindle (Fig. 1A) (Skoufias et al., 2006).

**Figure 1:**
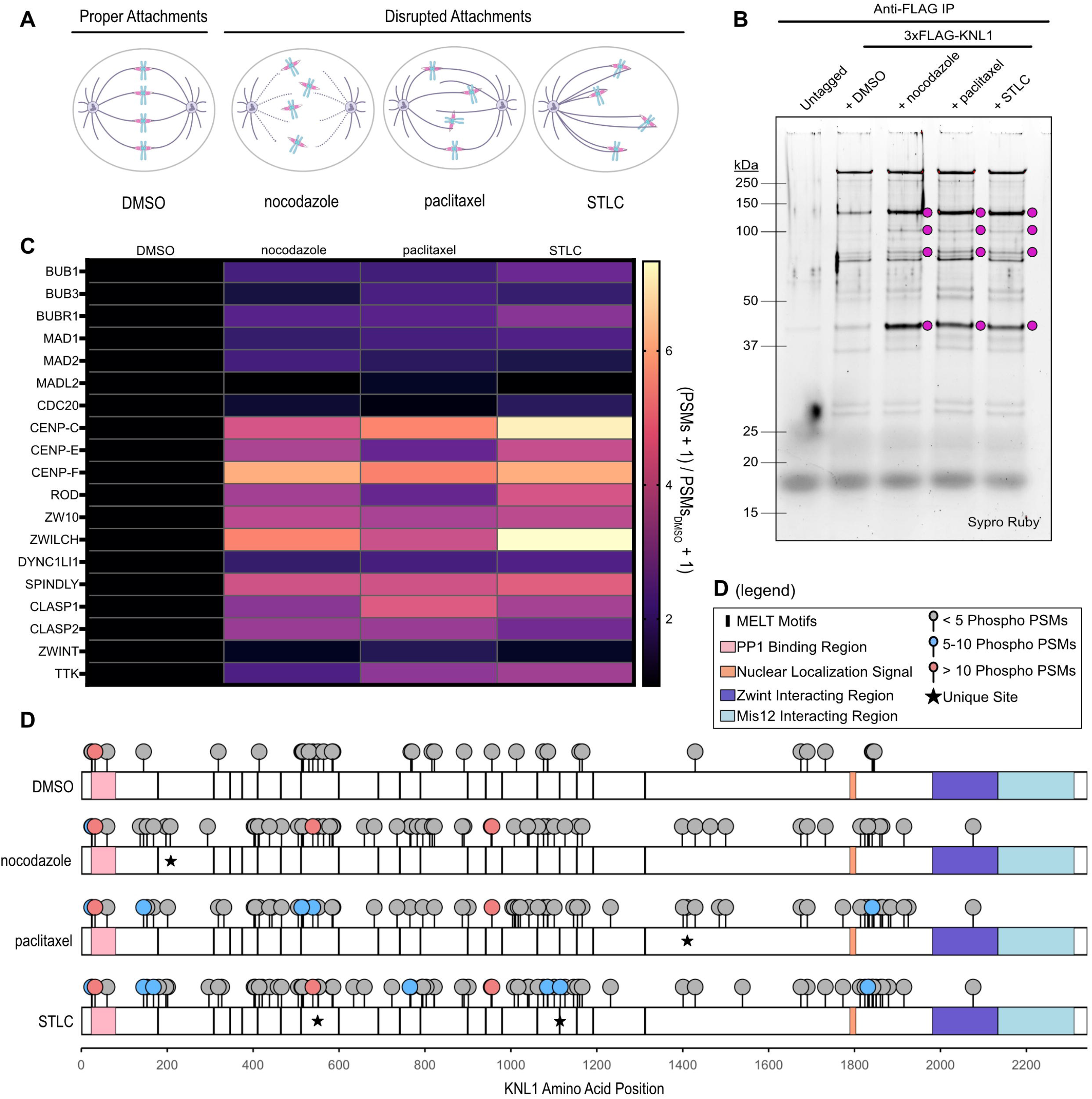
Human KNL1 is heavily phosphoregulated by kinetochore-microtubule attachment state. A. Schematic illustrating the effects of nocodazole, paclitaxel, and STLC on kinetochore-microtubule attachment state. The mitotic spindle is shown in purple, chromosomes are shown in blue, and kinetochores are shown in pink. B. 3xFLAG-KNL1 and interacting proteins purified from HEK 293T/17 cells by anti-FLAG immunoprecipitation. Cells were cultured in suspension to a density of 2.5e6 cells/mL, then treated with 300 nM nocodazole, 1 µM paclitaxel, 3 µM STLC, or an equivalent volume of DMSO. Cells were harvested 18 hours after the addition of compounds. Following purification, samples were visualized with Sypro Ruby protein stain after SDS-PAGE. Magenta circles represent bands that show enrichment in compound-treated cells. C. Liquid chromatography tandem mass-spectrometry (LC-MS/MS) proteomics analysis of spindle assembly checkpoint- and fibrous corona-associated proteins. Columns, from left to right: DMSO, nocodazole, paclitaxel, STLC. Brightness is proportional to the number of PSMs of a given kinetochore protein (on the vertical axis) relative to the number of PSMs present in the DMSO condition. Heatmap reflects average values across 3 replicates. The formula used to calculate these values was: [(average # of PSMs in ____ + 1)/(average # of PSMs in DMSO + 1)]. D. Phosphorylation profile of KNL1 under each treatment condition, plotted on a linear schematic. Rows, from top to bottom: DMSO, nocodazole, paclitaxel, STLC. Phosphorylation marks are color-coded by the average number of times a given event was detected across 3 mass spectrometry replicates (< 5 shown in gray, 5-10 shown in blue, > 10 shown in coral). Sites marked with a star are unique to one treatment condition and were detected in at least 2 out of 3 replicates.

To purify KNL1, we used CRISPR-Cas9 gene editing to introduce a 3xFLAG epitope tag at the endogenous KNL1 locus in HEK 293T/17 cells, then performed anti-FLAG immunoprecipitation on cells treated with each of the previously described microtubule-disrupting compounds, as well as an untagged, untreated HEK293T/17 control. SDS-PAGE analysis of anti-FLAG purifications revealed a discrete protein banding pattern compared to purifications from untagged cells (Fig. 1B). The banding patterns of 3xFLAG-KNL1 purifications were similar and highly reproducible. Additionally, we identified 4 proteins that migrate with apparent molecular weights of approximately 40, 75, 100, and 120 kDa that were more prominent following treatment with microtubule-disrupting compounds (Fig. 1B).

To gain a higher resolution interaction profile of KNL1 across attachment states, we performed liquid chromatography-tandem mass spectrometry (LC-MS/MS)-based proteomics (publicly available at https://massive.ucsd.edu/ProteoSAFe/dataset.jsp?task=61de21f1c11e4f8691829734e4f8855d). Across three independent replicates, we recovered high-confidence trypsin-digested peptides of KNL1 covering approximately 70-80% of the KNL1 amino acid sequence, with an average of 20-35 peptide spectrum matches (PSMs) per position (average coverage & depth of coverage: DMSO, 72%, 20.2; nocodazole, 73.57%, 30.8; paclitaxel, 73.67%, 30.1; STLC, 78.33%, 30.5) (Extended Data 1). In agreement with the SDS-PAGE results, mass spectrometry analysis confirmed that the interaction profile of KNL1 was similar across mitotically arrested cells, and detection of kinetochore proteins in untagged cells was minimal (Fig. 1C, Extended Data 2).

Given the importance of KNL1 phosphorylation in spindle assembly checkpoint activation and corona recruitment, we next extracted phosphorylation events from our proteomics dataset. Phosphorylated serine, threonine, and tyrosine residues from each experimental replicate were initially filtered for >75% confidence and ≥2 phosphorylated peptide spectrum matches (phospho-PSMs). The sum of sites that met these criteria, across 3 experimental replicates, was 35 in DMSO-, 67 in nocodazole-, 69 in paclitaxel-, and 79 in STLC-treated cells (Fig. 1D, Extended Data 3). Of these, 22 sites were shared across all 4 conditions, while 19 were shared only in mitotically arrested cells (Fig. 1D). We detected phosphorylation at 8 out of the 19 established MELT motifs, with increased MELT phosphorylation detected following compound treatment (DMSO, 1 MELT; nocodazole, 5 MELTs; paclitaxel, 5 MELTs; STLC, 7 MELTs) (Fig. 1D, Extended Data 3).

We then evaluated the frequency of phosphorylation at each site by calculating the average number of phospho-PSMs detected across three experimental replicates. The KNL1 residue with the highest average phospho-PSMs was S32 in all 4 conditions, representing ∼45-50% of the total peptides detected at this position (average phospho-PSM counts: DMSO, 17.00; nocodazole, 18.33; paclitaxel, 17.67; STLC, 19.00). Compared to asynchronous cells, we detected more phospho-PSMs at residues T539, S765, and S956 in all three mitotically arrested conditions (Extended Data 3). Comparison between nocodazole-, paclitaxel-, and STLC-treated cells revealed enrichment of phospho-S1831 in STLC-treated cells (average phospho-PSM counts: DMSO, 0; nocodazole, 0.67; paclitaxel, 2.00; STLC, 8.00). Phosphorylation at T1115 and T552 was unique to STLC treatment and detected in all 3 replicates. Phosphorylation at S207 was specific to nocodazole-treated cells, with 2 phospho-PSMs detected in 2 out of 3 replicates. T1410 phosphorylation was detected only in paclitaxel-treated cells and was found in 2 out of 3 experimental replicates, with 2 and 3 phospho-PSMs, respectively. Together, these data demonstrate that disruption of kinetochore-microtubule attachments induces frequent and widespread phosphorylation of KNL1.

In sum, we found that the binding partners of KNL1 were consistent across microtubule-disrupted attachment states, where KNL1 primarily bound outer kinetochore complexes Mis12 and Ndc80, in addition to spindle assembly checkpoint and fibrous corona-associated proteins. Compound treatment enhanced KNL1 interactions with several protein partners, which we hypothesize to be Ndc80 (75 kDa) and TTK (97 kDa), as well as the spindle assembly checkpoint-associated proteins Bub3 (37 kDa), Bub1 (120 kDa), and BubR1 (120 kDa) (Fig. 1B). Similarly, inner kinetochore proteins were more abundant in mitotically arrested cells, likely because inner and outer kinetochore association peaks during mitosis in mammalian cells. We did not identify compound-specific interactions, suggesting that there is a core set of proteins that respond to diverse kinetochore-microtubule attachment states.

In our mass spectrometry-based survey of KNL1 phosphorylation, we detected 111 phosphorylation sites on KNL1 across treatment groups, with 2-3 times more sites in compound-treated cells compared to DMSO-treated cells. At the residue level, S32 was the most heavily phosphorylated in all four conditions. Due to its abundance and consistency across treatment conditions, we speculate that phosphorylation of S32 may regulate a key function of KNL1.

Phosphorylation at S765 was enriched in mitotically arrested cells relative to DMSO-treated cells. Phosphorylation at this site has been previously reported (Ochoa et al., 2020), but the functional impact of KNL1 S765 phosphorylation has yet to be investigated. S765 is also within a region of KNL1 that was identified as a hit in a previous screen for essential protein domains (Herman et al., 2022). S765 lies within a consensus motif for Aurora B, the major kinase that regulates the error correction pathway (Welburn et al., 2010; Krenn and Musacchio 2015). Based on these data, we hypothesize that Aurora B phosphorylation of S765 may contribute to the functions of KNL1 in responding to low-tension kinetochore-microtubule attachments.

Mitotic enrichment was also evident for phospho-T539 and phospho-S956, sites which have been previously reported in a broad-scale phospho-proteomic analysis of mitotic cells (Olsen et al., 2010). S32, T539, and S956 are all followed by proline residues. Thus, we hypothesize that their putative kinase(s) are members of the proline-directed cyclin-dependent kinase (CDK) family, a class of kinases that regulate the cell cycle (Hayward et al., 2019; Pluta et al., 2023). While S32 phosphorylation was highly abundant across treatment conditions, phosphorylation at T539 and S956 was more prominent in mitotically arrested cells and, therefore, likely facilitated by CDK1, the primary mitotic CDK (Hayward et al., 2019).

We identified several potential attachment-specific sites, including T1115, T552, S1831, S207, and T1410. T1115 lies within a MELT sequence and was only detected in STLC-treated cells (McAinsh and Kops 2023). While this may reflect a difference in attachment-state regulation, it may also be attributed to higher coverage at this residue in STLC-treated cells. Phospho-S1831, a previously identified non-MELT TTK substrate (Olson et al., 2010), was present across compound-treated conditions but enriched in paclitaxel and STLC, relative to nocodazole. Additional novel, unique sites include T552 in STLC-, S207 in nocodazole-, and T1410 in paclitaxel-treated cells. For each of these sites, the average proportion of phosphorylated peptides was low; however, this could reflect that we are isolating a large pool of non-kinetochore bound KNL1 and some of these modifications are spatially regulated (Liu et al., 2009; Hayward et al., 2022). Additionally, our ability to detect these lowly-abundant sites likely reflects our deep proteomic coverage of KNL1.

This study provides an initial, semi-quantitative analysis of KNL1 phosphorylation following treatment with microtubule-disrupting compounds. Additional work is required to understand the functional importance of these phosphorylation events. Given the degeneracy of MELT motifs within KNL1, it is likely that no single site is uniquely essential to the spindle assembly checkpoint or error correction pathways; however, it will be interesting to understand how these phosphorylation events might provide additional regulation in the presence of low-tension kinetochore-microtubule attachments. Further, the experimental approaches we applied here may be expanded to investigate the phosphoregulation of other kinetochore proteins across attachment states.

## Methods

### Cell Culture

All experiments were conducted with HEK 293T/17 cells (ATCC CRL-11268). For adherent culture, cells were cultured in DMEM (Thermo 11-995-073) supplemented with 10% FBS and 2 mM L-glutamine and grown at 37 °C and 5% CO_2_ (complete DMEM). HEK 293T/17 cells were adapted to suspension culture by serial passaging in a combination of complete DMEM and FreeStyle™ 293 Expression Medium (Thermo 12-338-026). Adaptation occurred at 37 °C and 8% CO_2_ and cells were grown for 1 passage in 75% DMEM:25% FreeStyle; 1 passage in 50% DMEM:50% FreeStyle; 1 passage in 25% DMEM:75% FreeStyle; and 1 passage in 10% DMEM:90% FreeStyle. Following passage into 10% DMEM:90% FreeStyle, cells were trypsinized, washed 1x with FreeStyle media, and seeded as 25 mL cultures at ∼1-2e6 cells/mL in 100% FreeStyle media in 125 mL vented Erlenmeyer flasks (Celltreat 229821), shaking at 125 rpm. Suspension cells were monitored by visual inspection and counting and maintained at approximately 0.4e^6^ – 2e^6^ cells/mL with a flask:culture volume ratio of 4-5:1.

### 3xFLAG-KNL1 CRISPR Knock-in

The *KNL1* targeting crRNA and 3xFLAG-6xHis-KNL1 homology-directed repair (HDR) template were designed using the IDT Alt-R™ CRISPR HDR Design Tool. The 3xFLAG-6xHis-KNL1 HDR template was ordered as a gBlock, amplified by PCR, and purified using the Qiagen PCR Cleanup kit. CRISPR-Cas9 ribonucleoprotein (RNP) complexes were prepared according to the manufacturer’s recommendations. Briefly, *KNL1* targeting sgRNA was prepared by combining equal volumes of 100 μM crRNA and ATTO™ 550 tracrRNA (IDT 1075928), heating at 95 °C for 5 minutes, and cooling to room temperature. CRISPR-Cas9 RNP was prepared by combining 65 pmol of sgRNA (1.2 μL of 50 μM sgRNA) and 50 pmol (0.8 μL of 62 μM Cas9 (IDT 1081058)) in a 5 μL reaction volume supplemented with D-PBS (Thermo 14-190-250) and incubating for 30 min at 37 °C. CRISPR-Cas9 RNP and HDR template were combined (5 μL RNP + 5 μL HDR template at ∼200 ng/μL) and delivered to HEK 293T/17 cells using the SF Cell Line 4D-Nucleofector® X Kit (Lonza V4XC-2032) according to the manufacturer’s recommendations. Cells were collected by trypsinization, centrifuged for 5 minutes at 300x *g*, and resuspended in fresh media. Cells were counted and approximately 200,000 cells/nucleofection were pelleted by centrifugation for 10 min at 90x *g*. Cells were gently resuspended in 20 μL Buffer SF + supplement/200,000 cells, mixed with CRISPR-Cas9 RNP + HDR template, and loaded into nucleofector strip cassettes. Cells were electroporated with program CM-130 on a 4D-Nucleofector® X Unit (Lonza AAF-1003B and AAF-1003X). Following nucleofection, cells were transferred to 96-well plates containing prewarmed recovery medium (complete DMEM supplemented with 1 μM AZD7648, and 3 μM ART558) and incubated at 37 °C. Cells were plated at low density 3-4 days post-nucleofection and individual subclones were isolated and expanded. Cells were screened by PCR to identify targeted clones, and multiple clones with evidence of homozygous targeting were expanded.

### Treatment

Cells were expanded to 250 mL suspension cultures and treated with compounds at a density of ∼2.5e6 cells/mL. Each flask received either 300 nM nocodazole, 1 μM paclitaxel, 3 μM *S*-trityl-L-cysteine, or an equivalent volume of DMSO. Cells were incubated for 18 hours following treatment.

### Cell harvest

Cells were harvested according to a protocol similar to that described in (LaCava et al., 2016) Cells were pelleted via centrifugation at 1,000x *g* for 10 minutes at 4 °C. Cell pellets were washed two times by resuspending cells in 50 mL of cold PBS, followed by centrifugation at 1,000x *g* for 5 minutes at room temp. After washing, cell pellets were resuspended in 10 mL of cold PBS and pipetted into capped 10 mL syringes. Syringes with cell suspensions were placed into 50 mL conical tubes and centrifuged at 1,000x *g* for 5 minutes to pellet cells in the syringe. Syringes were removed from conical tubes, PBS was aspirated, and cells were dripped via syringe into 50 mL conical tubes filled with liquid nitrogen. Small holes were made on tube lids, and excess nitrogen was removed by inverting capped tubes. Frozen cell droplets were then pulverized by cryogenic milling using a Spex 6875 Freezer/Mill. The duty cycle was 2 min ‘ON’, 2 min ‘OFF’ at a rate of 10 Hz; 10 cycles were performed.

### Anti-FLAG bead preparation

Protein G magnetic beads (Thermo 100.09D) were washed twice with 1 mL of 0.1 M Na-Phosphate buffer. Beads were then incubated with anti-FLAG antibody at a concentration of 0.25 μg antibody per 1 μL of beads for 30 min at room temperature with gentle agitation. Following antibody binding, beads were washed twice with 1 mL of 0.1 M Na-Phosphate + 0.01% Tween 20, then two more times with 1 mL of 0.2 M triethanolamine. Following these washes, beads were resuspended in 1 mL of 20 mM dimethyl pimelimidate (DMP; Sigma D8388) in 0.2 M triethanolamine and incubated with end-over-end rotation for 30 minutes at room temp. The crosslinking reaction was quenched by the removal of DMP and incubation with 1 mL of 50 mM Tris-HCl, pH 7.5 for 15 minutes with end-over-end rotation at room temp. Beads were washed 3 times with PBS + 0.1% Tween 20 (PBS-T) and resuspended in PBS-T at a volume equivalent to the starting volume of beads. Beads were stored at 4 °C and used within 1 week of preparation.

### Immunoprecipitation

For each sample, 2 grams of pulverized cell powder was weighed out on liquid nitrogen. Cell lysates were prepared by resuspending cell powder in Buffer H 0.15 (25 mM HEPES, pH 8.0; 150 mM KCl; 2 mM MgCl_2_; 0.1 mM EDTA, pH 8.0; 0.5 mM EGTA-KOH, 15% glycerol, 0.1% NP-40) on ice at a ratio of 2 mL Buffer H 0.15:1 g of cell powder. Lysis buffer was supplemented with 0.4 U/μL Benzonase; phosphatase inhibitors (0.1 mM sodium orthovanadate, 2 mM β-glycerophosphate, 1 mM sodium pyrophosphate, 5 mM sodium fluoride, and 0.2 μM microcystin-LR); and protease inhibitors (200 μM phenylmethylsulfonyl fluoride, 100 μg/mL leupeptin, 100 µg/ml pepstatin A, and 100 µg/ml chymostatin). Lysates were clarified by centrifugation at 21,000x *g* for 10 min at 4 °C. Clarified lysates were incubated with anti-FLAG beads (prepared as described above; 60 μL beads/1g of cell powder) for 30 minutes at 4 °C with end-over-end mixing. Beads were recovered by magnetization and washed 5 times with 1 mL of wash buffer (Buffer H 0.15 supplemented with protease and phosphatase inhibitors as above). Bead-bound proteins were eluted via incubation with 1 mg/mL 3xFLAG peptide in Buffer H 0.15 + protease inhibitors for 40 minutes with gentle agitation at room temperature.

### Mass-Spectrometry

Ten (10) µL of each elution was analyzed by SDS-PAGE and Sypro Ruby staining. To perform mass spectrometry, we removed excess 3xFLAG peptide by running the remainder of each sample into a 4-12% Bis-Tris gel for 5 minutes at 100 V such that all proteins with a molecular weight greater than 3 kDa remained unseparated. The gel was stained overnight with 0.1% Coomassie G-250 and destained the following day until destain solution ran clear. The gel was then rinsed with deionized water for 10 minutes. For each immunoprecipitation, the band containing all proteins above 3 kDa was cut out of the gel. The gel plug for each sample then underwent trypsin digestion as previously reported (Wilm et al., 1996).

LC-MS/MS analysis was performed by an Orbitrap Eclipse Tribrid mass spectrometer (Thermo Scientific) operated in positive ion mode with a FAIMS interface. The Easy-nLC 1000 (Thermo Scientific) LC system consisted of a fused-silica nanospray needle (PicoTip™ emitter, 50 µm ID x 20 cm, New Objective) packed in-house to 27 cm with ReproSil-Pur 120 C18-AQ (Dr. Maisch) and a trap (IntegraFrit™ Capillary, 100 µm ID x 2 cm, New Objective) packed in-house to 2 cm with Magic C18-AQ (Michrom) with mobile phases of 0.1% formic acid (FA) in water (A) and 0.1% FA in MeCN (B). The peptide sample was diluted in 20 µL of 0.1% FA, 2% MeCN and 18 µL was loaded onto the column and separated over a total run time of 115 minutes at a flow rate of 300 nL/min with a gradient from 2 to 8% B for 3 min, 8 to 30% B for 87 min, 30 to 45% B for 10 min, 45 to 60% B for 3 min, 60 to 95% B for 2 min, hold 95% B for 11 min. FAIMS MS/MS analysis occurred over a 3 second cycle time consisting of 1 full scan MS from 375-1500 m/z at resolution 240,000 followed by data dependent MS/MS scans cycling through FAIMS CV of -40, -60 and -80 using HCD activation with 27% normalized collision energy of the most abundant ions. Selected ions were dynamically excluded for 60 seconds after a repeat count of 1.

The MS/MS results were searched using Proteome Discoverer v3.1 against a human Universal Protein Resource (UniProt) sequence database downloaded on 11/1/2023 containing common contaminants with tryptic enzyme constraint set for up to two missed cleavages, oxidized methionine and phosphorylation of serine, threonine and tyrosine set as a variable modification, and carbamidomethylated cysteine set as a static modification. Peptide MH+ mass tolerances were set at 10 ppm. The results were filtered to include high-confident (peptide FDR < 1%) identifications.

PSM, phosphorylation, and coverage data was analyzed in Excel. For data shown in Figure 1D, fold change in PSMs for a given kinetochore protein was determined by the raw number of PSMs for a compound-treated condition divided by the raw number of PSMs for the DMSO-treated condition. 1 PSM was added to all raw RSM counts to avoid division by 0. Fold change in PSMs was then plotted in GraphPad Prism. Phosphorylation schematics and coverage maps were made with ggplot2 in R.

## Reagents

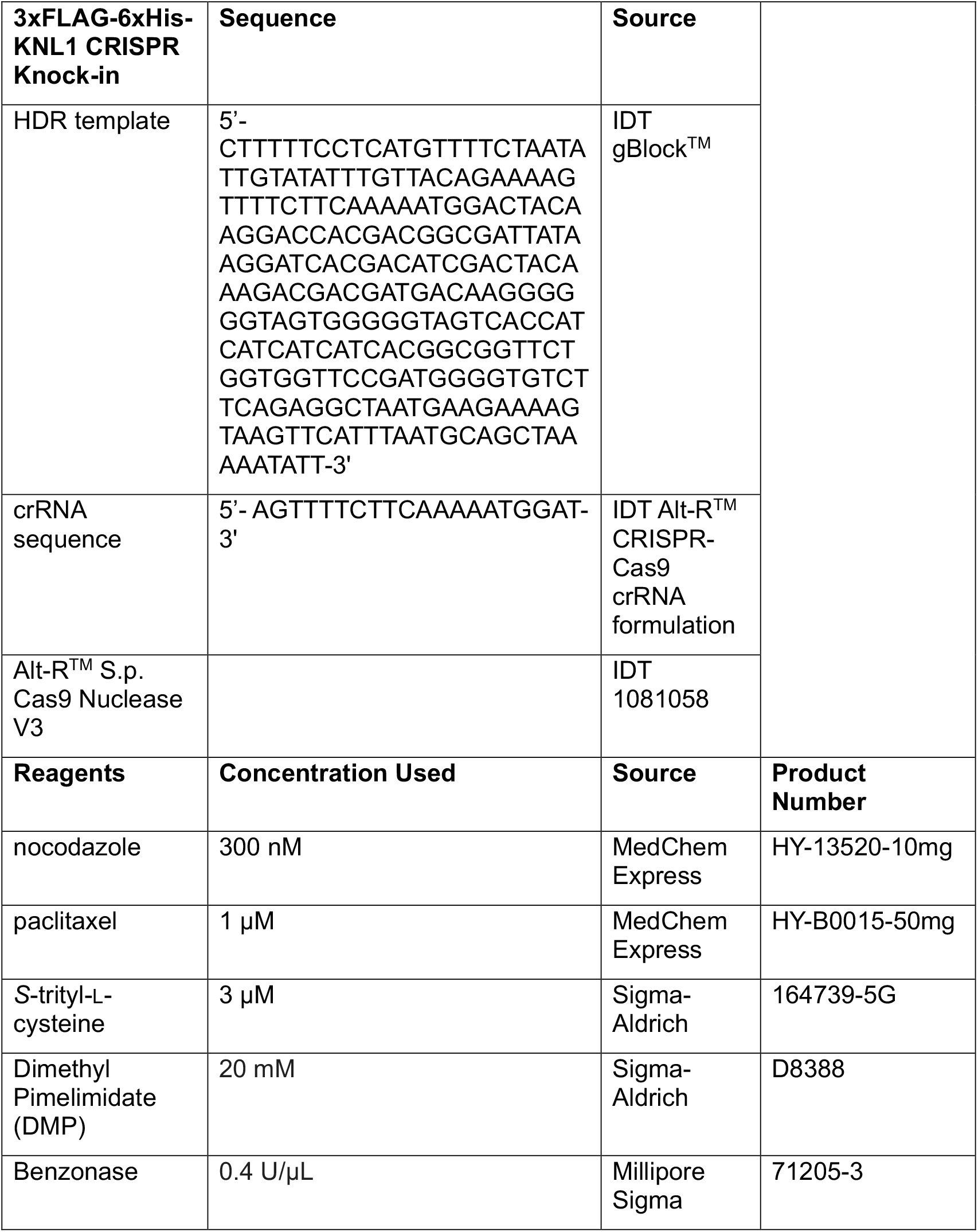
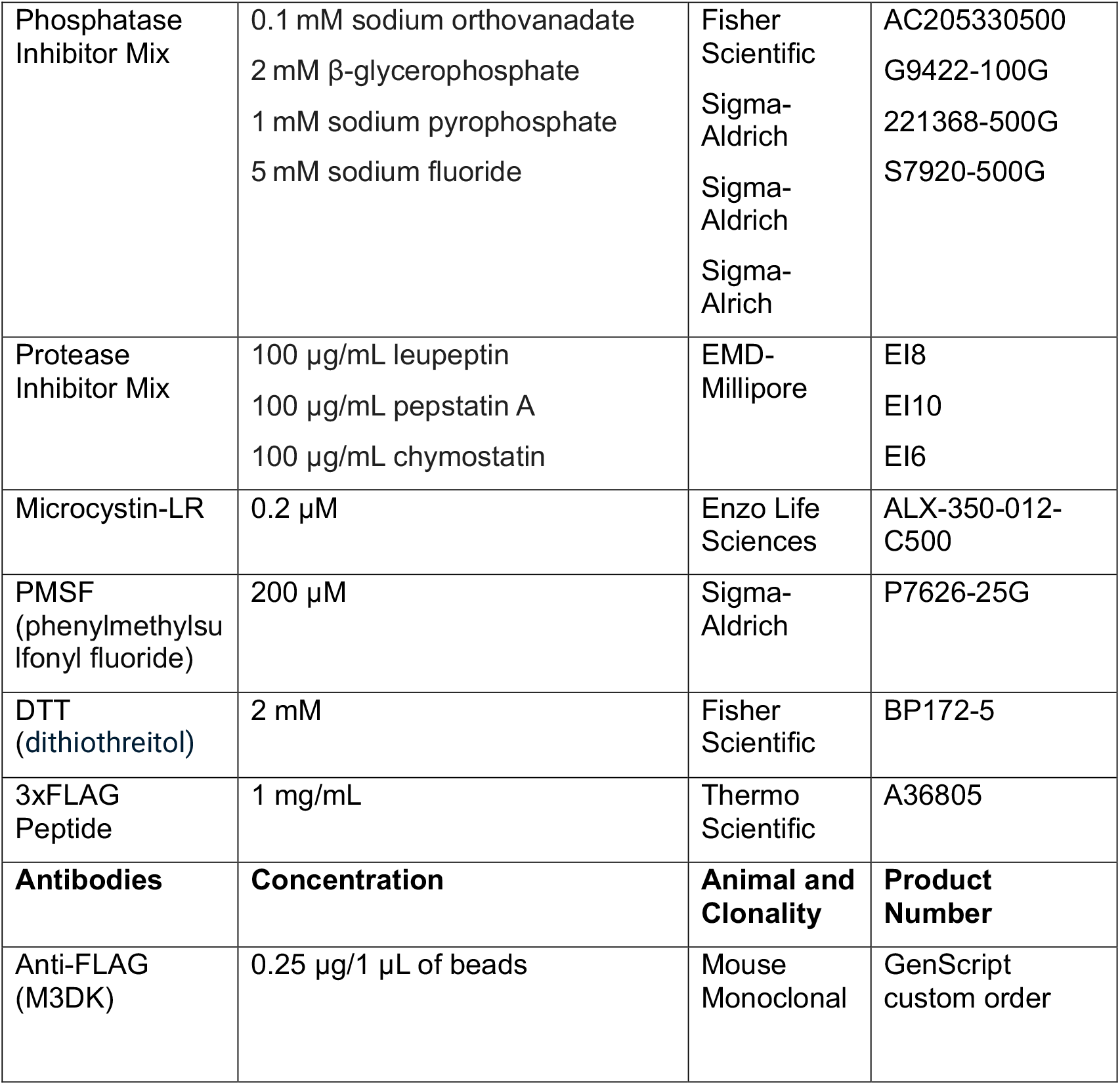

## Extended Data

<u>Extended Data 1</u> Coverage maps of KNL1 illustrating average coverage for each condition across 3 mass-spectrometry experiments. Maps are aligned with corresponding phosphorylation schematics from Fig. 1D

<u>Extended Data 2</u> PSM data for all kinetochore proteins, in all treatment conditions, for the mass spectrometry results shown in Fig. 1C

<u>Extended Data 3</u> KNL1 phosphorylation and average coverage at phospho-sites

https://massive.ucsd.edu/ProteoSAFe/dataset.jsp?task=61de21f1c11e4f8691829734e4f8855d is a link to all mass spectrometry data

## Acknowledgements

We would like to thank the members of the Biggins lab for their insightful feedback and advice throughout this study, as well as the Proteomics and Metabolomics shared resource at Fred Hutch Cancer Center for mass spectrometry sample processing, data collection, and analysis.

## Funding

This research was supported by NIH P30 CA015704 of the Fred Hutch/University of Washington/Seattle Children’s Cancer Consortium, which includes the Proteomics and Metabolomics Shared Resource, RRID:SCR_022618.

This work was supported by NIH R35 GM149357 to SB and NIH F32 GM156071 to ARF.

SB is an Investigator of the Howard Hughes Medical Institute.

## Author Contributions

Abby Jurasin: Data Curation, Formal Analysis, Investigation, Validation, Visualization, Writing – original draft

Anderson Frank: Conceptualization, Formal Analysis, Funding acquisition, Investigation, Methodology, Supervision, Writing – review & editing

Sue Biggins: Conceptualization, Formal Analysis, Funding acquisition, Project administration, Supervision, Writing – review & editing

