## Supplementary figures and images for "Proteomics reveals extensive phosphoregulation of outer kinetochore protein KNL1"

### supplementary data 1

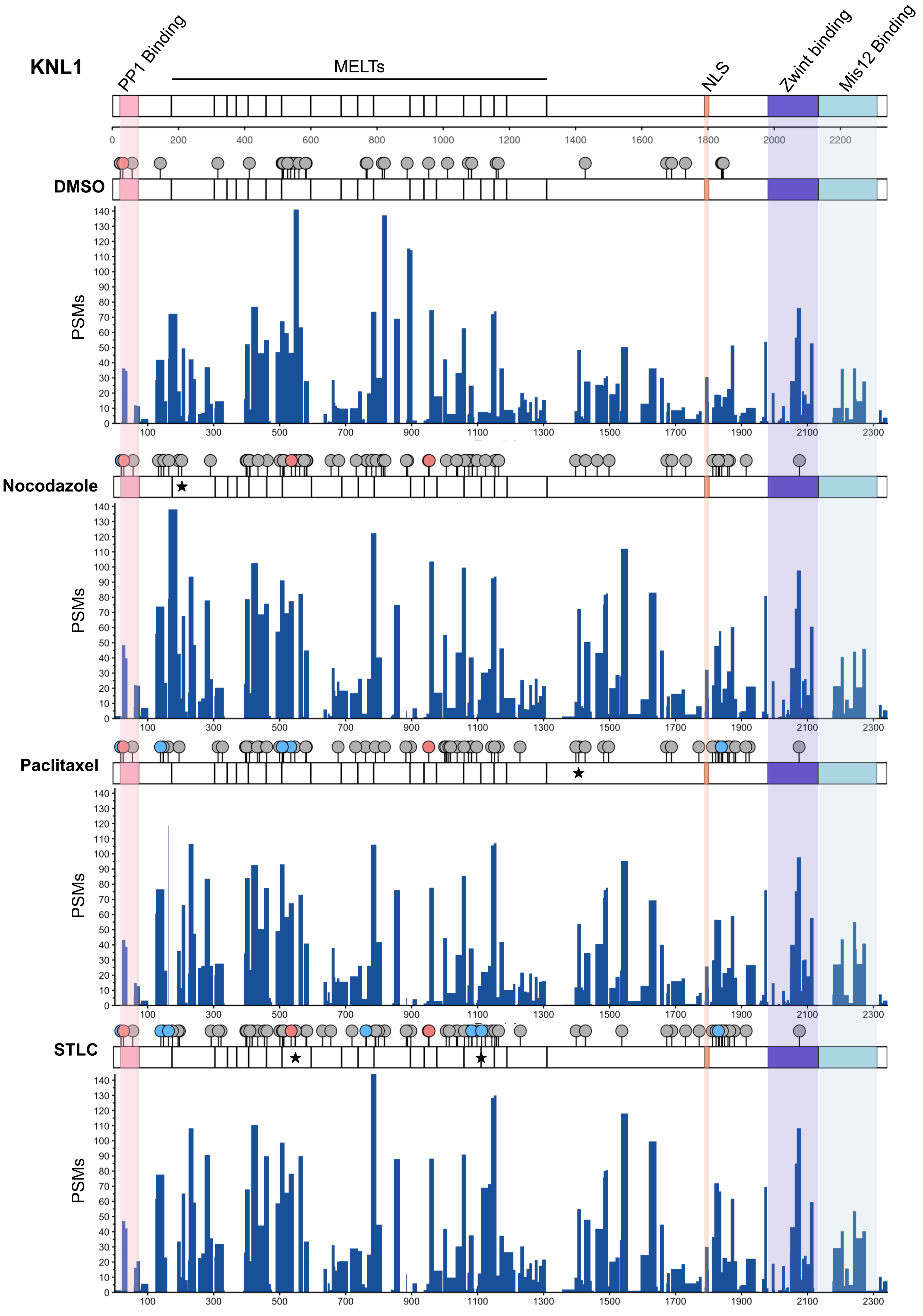
